# Spatiotemporal dynamics and emergent ordering in a mixture of morphologically distinct bacteria having different cell motility

**DOI:** 10.1101/2023.07.13.548839

**Authors:** Kaustav Mondal, Palash Bera, Pushpita Ghosh

## Abstract

Microbial communities exhibit complex behaviors driven by species interactions and individual characteristics. In this study, we delve into the dynamics of a mixed bacterial population comprising two distinct species with different morphology and motility aspects. Employing agent-based modeling and computer simulations, we analyze the impacts of size ratios and packing fractions on dispersal patterns, aggregate formation, clustering, and spatial ordering. Notably, we find that motility and anisotropy of elongated bacteria significantly influence the distribution and spatial organization of nonmotile spherical species. Passive spherical cells display superdiffusive behavior, particularly at smaller size ratios, while active rod-like cells exhibit normal diffusive behavior in the diffusion regime. As the size ratio increases, clustering of passive cells is observed, accompanied by enhanced alignment and closer packing of active cells in the presence of higher passive cell area fractions. As the size ratio increases, clustering of passive cells is observed, accompanied by enhanced alignment and closer packing of active cells in the presence of higher passive cell area fractions. Additionally, we identify the pivotal role of passive cell area fraction in influencing the response of active cells toward nematicity, with its dependence on size ratio. These findings shed light on the significance of morphology and motility in shaping the collective behavior of microbial communities, providing valuable insights into complex microbial behaviors with implications for ecology, biotechnology, and bioengineering.

## Introduction

In recent years, the study of active systems composed of self-propelled units has emerged as a captivating field of research, offering unique insights into non-equilibrium dynamics.^1^ Active systems exhibit collective behavior across various length scales, spanning from microscopic levels such as bacterial colonies,^2^ cell cytoskeletons,^3^ and active nematics,^4–6^ to macroscopic scales encompassing bird flocks, ^7^ human crowds,^8^ and animal herds.^9^ This diverse range of active systems has led to the observation of intriguing phenomena, including self-regulation,^1^ swimming,^10^ nonequilibrium disorder-order transitions,^11^ pattern formation,^12^ clustering,^13–17^ segregation, and rheological phase behavior.^18–20^ Among the distinctive dynamic phenomena exhibited by active systems, motility-induced phase separation (MIPS) stands out prominently. MIPS refers to the phenomenon where active particles undergo spontaneous separation, forming distinct solid-gas phases. This intriguing behavior has been experimentally validated through a number of studies and has also been investigated numerically to gain a deeper understanding of its underlying mechanisms.^16, 21–24^

The interactions between active and passive particles have recently gained significant attention due to their characteristic dynamics. Passive colloids, influenced by Brownian motion, have the ability to self-organize and exhibit a diverse range of phases depending on their morphologies and the interplay of particle-particle contact forces. In active media, cluster aggregation occurs as a result of a net attractive force between colloidal particles. Notably, Wu et al.^25^ conducted experiments, providing the first evidence of passive colloid motion within an active suspension. Theoretical studies on active-passive mixtures have predicted segregation phenomena based on variations in motility^24, 26, 27^ or diffusivity.^28, 29^ These studies contribute to our understanding of active-passive particle interactions and hold potential for uncovering new phenomena and applications across various fields.

In a recent experiment,^30^ *E. coli* bacteria were utilized as active cells, demonstrating the self-assembly and diverse phase behavior of colloid particles in active conditions. It was observed that the relative size of the colloids played a significant role in clustering. Similarly in another experiment,^31^ using *P. aurantiaca* cells as active agents and silica as colloids significant clustering of colloids at high densities of active cells was reported. Motivated by these experimental findings, our study aims to investigate the clustering behavior of nonmotile spherical passive cells in the presence of motile, rod-shaped active cells using an agent-based model and computer simulations. Previous research on self-propelled hard rods has revealed phenomena such as flocking and alignment through collisions, where increasing the length of the active cells leads to stronger alignment and group formation.^32–34^ Additionally, studies by Sudha et al. ^35^ have shown that mutual attraction between colloidal particles in an anisotropic fluid, like a nematic liquid crystal phase, can result in the formation of hierarchical aggregates. However, limited research has focused on how the aggregation of rod-shaped cells impacts the spatiotemporal dynamics of active-passive systems. Therefore, our study aims to explore the influence of the morphology and activity of rod-shaped active bacterial cells on the overall dynamics of the mixed system and investigate the role of cell size and motility in the phase behavior of morphologically distinct colloid-like passive particles. In our model, we consider two types of cells: non-motile spherical passive cells and motile elongated active cells. The key feature of the active cells is their self-propulsion or motility, while both types of cells interact mechanically through repulsive forces. The length of the elongated bacteria, denoted as l*_a_*, is a crucial parameter in our system, and we normalize it using the diameter of the passive cell, denoted as d*_p_*, to obtain the size ratio (S). Two other parameters we consider are the area fraction of passive cells (ϕ*_p_*) and the area fraction of active cells (ϕ*_a_*), which are varied during the study. Using an individual-based model, we conduct simulations and analyses to examine the phase behavior of both active and passive cells, how it depends on the size ratio, and how it influences the spatiotemporal dynamics of the mixture. Our results reveal that as the size ratio increases, the active cells undergo a nematic transition from an isotropic to an ordered state. The mean cluster size of passive cells increases with the size ratio (S). Furthermore, we observe that nonmotile cells exhibit higher phase separation and form larger clusters compared to motile cells. By utilizing the mean squared displacement (MSD) as a measure of spatial motion, we demonstrate that smaller size ratios lead to superdiffusive behavior in passive cells which is an interesting outcome of our study.

Overall, our theoretical study provides insights into the clustering and phase behavior of active-passive particles with an example of living system such as bacteria. By systematically varying the size ratio, we elucidate the interplay between cell characteristics and the spatiotemporal dynamics of the mixture. These findings contribute to our understanding of collective phenomena in active-passive systems and have implications for various fields, including biophysics and materials science.

## Model and Methods

We consider an individual-based model of active or motile rod-shaped and passive or nonmotile spherical bacterial cells in a two-dimensional box. Each active cell is represented by spherocylindrical particle with a fixed diameter d*_a_* and of variable length l*_a_* = l + d*_a_*, where l represents the particle’s cylindrical length. The spherical cells have fixed diameter of d*_p_*. Our system is a mixture of two types of bacterial cells: N_1_ number of active cells and N_2_ number of passive cells. An individual cell’s position and orientation are given by a spatial coordinate ⃗r(*x, y*) and a unit vector ⃗u(u*_x_*, u*_y_*). Each particle experiences a random force ^ζ⃗^ from the surrounding medium attributing the presence of a little randomness in their motion. The equation of motions for the active and passive cells follow an over-damped dynamics justifying a densely packed mixed-species colony where viscosity dominates over inertia. We assume that the particles are submerged in a viscous medium such that the linear velocity and the angular velocity are proportional to the force and torque acted upon particles; the equation of translational motion for the particles is given by the:

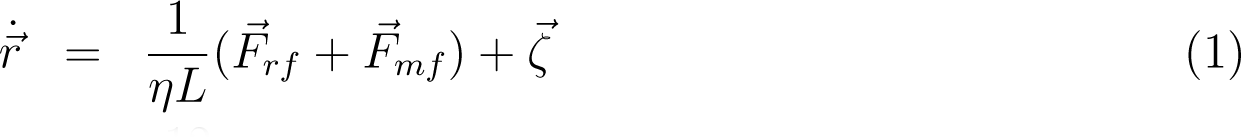

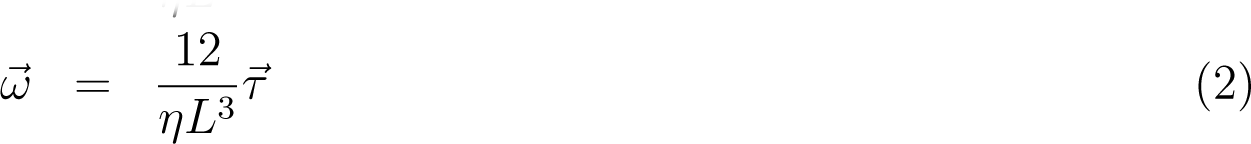

⃗where ⃗ṙ and ω are linear and angular velocities respectively and η represents the friction coefficient. In case of nonmotile spherical bacteria, the motility force component ^F⃗^*_mf_* = 0. Furthermore, each particle is subjected to a random force ^ζ⃗^ from the surrounding medium. It is drawn from a uniform distribution spanning the range −10*^−^*^3^ pascal.µm^2^ to +10*^−^*^3^ pascal.µm^2^. From eqn 1, we see that each active bacterial cell is equipped with a motility force of ^F⃗^*_mf_*, resulting in a self-propelled velocity acting along its long axis. Besides, each cell interact with its neighboring cells with a short-range repulsive mechanical force, ^F⃗^*_rf_* . During the time evolution of the system, we systematically corrected the center of mass velocity of the simulation box, thereby ensuring there is no external force acting on the system. According to the Hertzian theory of elastic contact, mechanical interaction between the cells follows short-range repulsive force which is modeled in prior studies ^36, 37^ as: ^F⃗^*_rf_* = Ed^1^*^/^*^2^h^3^*^/^*^2^. Here E is the elastic modulus of the cells and h = d_0_ − r represents the overlap between two interacting cells, where r corresponds to the closest distance of the approach between the two cells and d_0_ is the diameter is the cell Apart from the cell-cell repulsive mechanical interaction, the essential feature of an active cell is the presence of self-propulsive or motility force (^F⃗^*_mf_*). We considered this motility force acting along the orientation of the active cell. The generic functional forms of the motility force expression employed for our current work can be expressed as where f*_mot_* is the proportionality constant. We simulate the system in a square box of size (200 × 200)µm^2^ with periodic boundary conditions. The particles’s new positions and velocities are determined by solving the equations of motion using the simple Euler method with time step δt = 10*^−^*^3^s. We first initialize the system by seeding the active spherocylindrical particles randomly placed inside the simulation box. After some time, when the system equilibrates, the passive spherical particles are introduced into the system randomly. Following this initialization protocol, ensures that the modeled cells arrange naturally in a densely packed manner in a two-dimensional surface.

**Table 1:** Parameters, constants and expression of repulsive forces used in simulations

| Parametes | Symbols | Simulations |
| --- | --- | --- |
| Boxsize | $L_x \times L_y$ | $200 \times 200 \mu m^2$ |
| Active cell area fraction | $\phi_a$ | 0.3, 0.5 |
| Passive cell area fraction | $\phi_p$ | 0.1-0.5 |
| Diameter of active cell | $d_a$ | $1 \mu m$ |
| Diameter of passive cell | $d_p$ | $2 \mu m$ |
| Size ration | $S$ | 1.5-5.5 |
| Total time | T | 1600 s |
| time step | $\delta t$ | $10^{-3}$ s |
| motiltiy force constant | $f_{mot}$ | $5 \times 10^4$ pascal.$\mu m$ |
| friction constant | $\eta$ | 200 pascal.s |
| Elastic modulus (Cells) | E | $2 \times 10^5$ pascal |
| **Component pair** | **Repulsive forces** (pascal.$\mu m^2$) | |
| active-active | $E d_a^{1/2} (d_a - r)^{3/2}$ | |
| passive-passive | $E d_p^{1/2} (d_p - r)^{3/2}$ | |
| active-passive | $E (0.5 d_p + 0.5 d_a)^{1/2} ((0.5 d_p + 0.5 d_a) - r)^{3/2}$ | |

## Results and discussion

Our study focuses on the spatiotemporal dynamics and emergent ordering in a mixture of two bacterial species having different morphology and motility aspects. How the motility and anisotropy of one of the species can induce dispersal, aggregate formation and clustering of spherical nonmotile species is one of the significant issues that we have investigated through computer simulations and systematic analysis. We mainly emphasize on two key elements to study the spatiotemporal dynamics and organization of the two bacterial species: by varying surface fractions and by varying size-ratio between the two components. The surface or packing fraction is defined by ϕ = *<u>^Na^</u>* (where a is the area of a single cell, A is the total area of the box and N corresponds to the total number of cells in the simulation box) and the size ratio corresponds to S = *<u>^la^</u> p* (where l*_a_* corresponds to the length of the active cell and d*_p_* is the diameter of the passive cell).

### Variation of mean speed with size ratio and area fraction

We first examined the mean speed of active cells in relation to the size ratio (S) and packing fraction (ϕ). Figure 1(a) demonstrates the variation of the mean speed of the active cells with respect to the size ratio for three different packing fractions of passive cells keeping the active packing fraction at ϕ*_a_* = 0.3. Our results indicate that the mean speed of active cells decreases as the size ratio increases. This decrease can be attributed to the motility aspect of elongated cells. As the length of the active cells increases and the size ratio increases, the self-propulsion velocity of active cells diminishes at the individual level due to increased drag forces experienced by longer cells. This observation suggests that the size ratio has a significant impact on the motility of active cells within the bacterial mixture. Furthermore, we explored the variation of mean speed of active cell with the size ratio at a fixed surface fraction of passive cells (ϕ*_p_* = 0.3) for two different packing fractions of active cells(ϕ*_a_* = 0.3, 0.5), as shown in Figure 1(b). Our findings support our earlier observation, indicating that mean speed decreases with an increasing size ratio. However, we noted that higher packing fractions of active cells result in higher mean speeds, suggesting that the overall concentration of active cells plays a significant role in determinining their collective motility.

**Figure 1:**
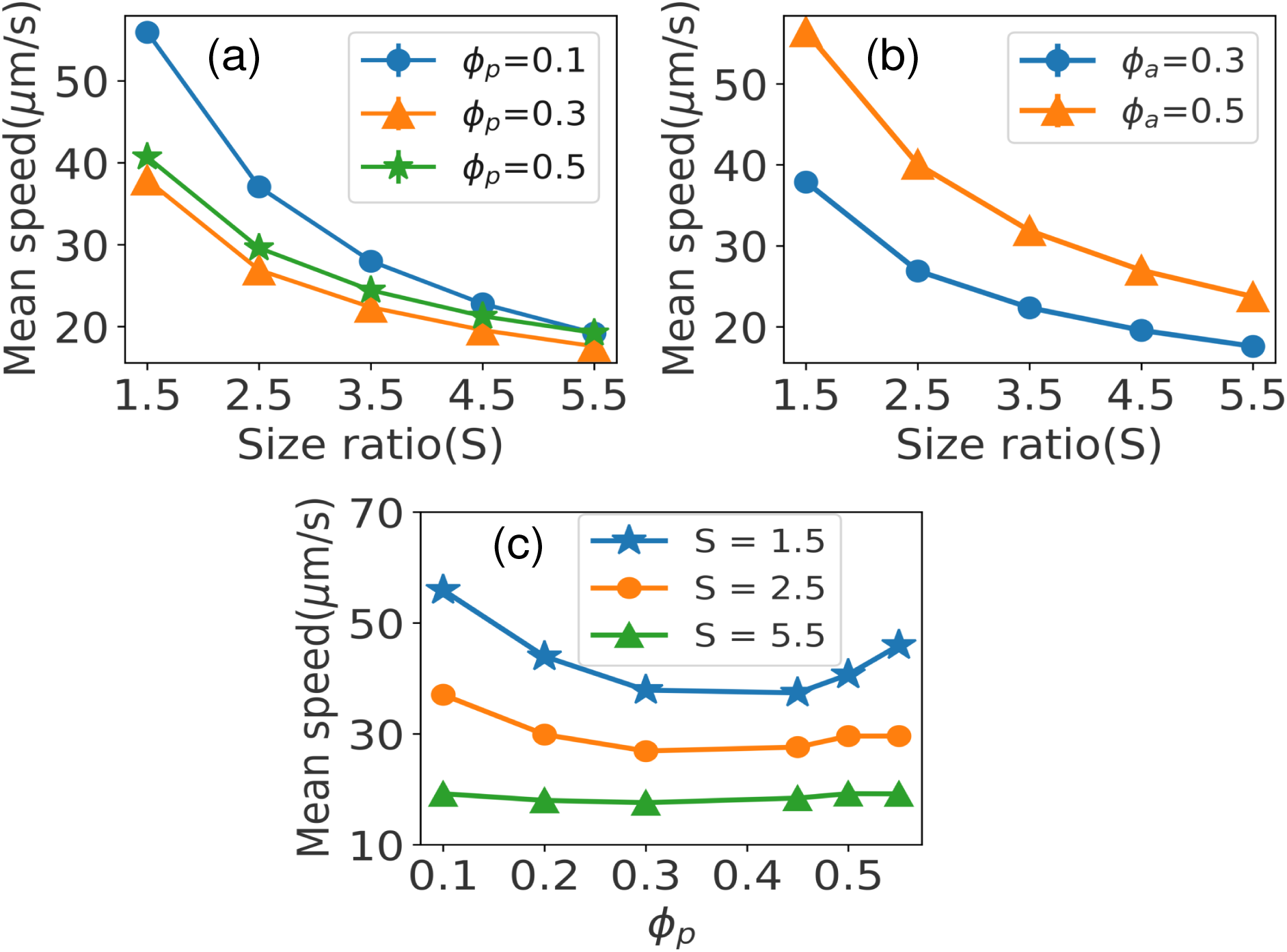
(a) Plot of mean speed of active cells as a function of size ratios (S) for different ϕ*_p_* and fixed ϕ*_a_* = 0.3 (b) Plot of mean speed of active cells as a function of size ratios (S) for different ϕ*_a_* and fixed ϕ*_p_* = 0.3 and (c) Plot of the mean speed of active cells as a function of ϕ*_p_* for different size ratio(S). Standard errors are equal to the marker size of the plot.

However, an interesting observation arises when examining Figure 1(a) for a fixed S. The result is shown in Figure 1(c) for three size ratios: S = 1.5, 2.5, and 5.5. For lower S(1.5, 2.5), when ϕ*_p_* is increased from 0.1 to 0.3, the mean speed of active cells decreases. This decrease can be attributed to the increased density of passive cells, which can impede the movement of active cells. Surprisingly, when ϕ*_p_* is further increased from 0.3 to 0.5, the mean speed of active cells shows an increase. This phenomenon could be explained by considering the interplay between the active and passive cells. At low passive packing fractions (e.g., ϕ*_p_* = 0.1), the environment may be relatively sparse, with minimal interactions among passive and active cells. As ϕ*_p_* increases to 0.3, the density of passive cells rises, leading to more frequent interactions and hindrance of active cells’ motion. However, when ϕ*_p_* reaches 0.5, the system may undergo a transition where the increased density of passive cells starts to facilitate the emergence of collective behavior. The interactions among passive cells and their coupling with active cells can lead to cooperative effects that leads to aggregation due to depletion-attraction allowing active cells to move more efficiently. This entropy-driven attraction becomes effective when the concentration and size of the depletant is high. In contrast, for higher size ratio (S = 5.5), this variation is insignificant suggesting that the system’s temporal dynamics remains unperturbed by changing the surface fraction of passive cells.

To gain further insights into the collective dynamics as observed from Figure 1, we estimated the average translational force experienced by each cell. This force provides valuable information regarding the underlying causes of the observed behavior. Here, we computed the total force per unit length of the active cells and analyzed its distribution. The distribution of the forces on active cells is shown in Figure 2(a) and (b), for three different S. The distribution of forces sheds light on the interplay between cell-cell interactions, motility, and the emergent ordering observed in the system. We observe that the net driving force on shorter cells is higher than on longer ones which aligns with the findings from our simulation videos-(Movie-1 -Movie-9) in Supporting Information (SI). As the size of active cells increases, they encounter more obstacles in the form of other active and passive cells, which hinders their self-propulsive motion. This phenomenon explains why the mean speed of active cells decreases with increasing size ratio.

**Figure 2:**
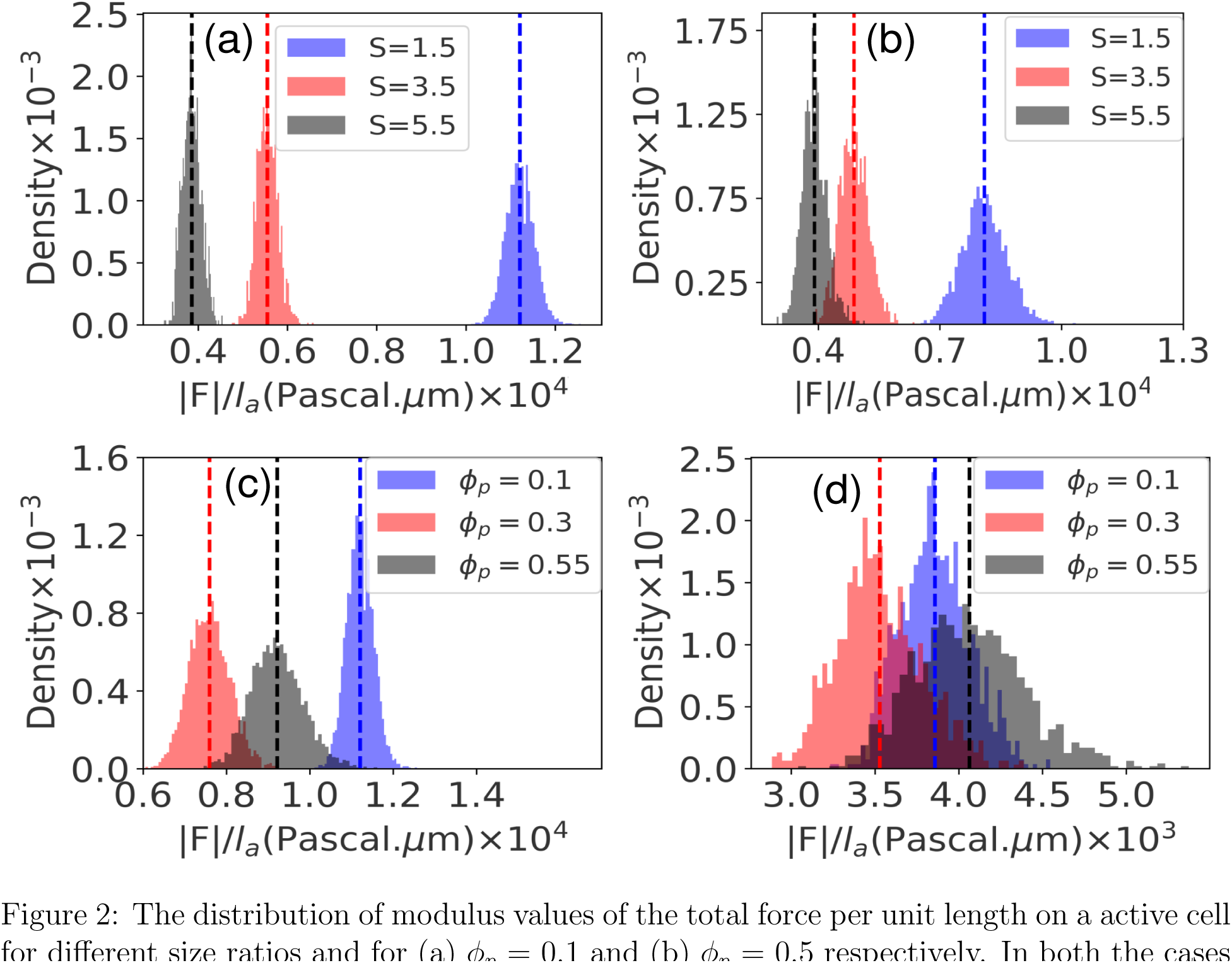
The distribution of modulus values of the total force per unit length on a active cell for different size ratios and for (a) ϕ*_p_* = 0.1 and (b) ϕ*_p_* = 0.5 respectively. In both the cases active cell packing fraction was kept fixed at ϕ*_a_* = 0.3. (c) and (d) shows the distribution of modulus values of the total force per unit length on a active cell for fixed S = 1.5 (smaller) and S = 5.5 (larger) respectively and ϕ*_a_* = 0.3. The dashed lines shows the mean of the corresponding distributions.

Another notable observation in our study is the effect of packing fraction (ϕ*_p_*) on the total translational force for a fixed size ratio as elucidated in Figure 2(c) and (d). For a fixed ϕ*_a_* = 0.3, we found that as the packing fraction increases, the total translational force initially decreases until reaching a critical value at around ϕ*_p_* ≈ 0.35. Beyond this point, the total translational force starts to increase again. Analysis of the simulation videos revealed that the decrease in total translational force at lower packing fractions is primarily attributed to increased collision events experienced by the active cells. These collisions hinder their motion, resulting in reduced mean speed. However, when the packing fraction exceeds ϕ*_p_* > 0.45, the abundance of passive cells leads to increased alignment and collective motion of the active cells. Consequently, the active cells exhibit more consistent forward motion, resulting in an overall increase in mean speed. Additionally, we noticed that the impact of packing fraction on the mean speed of active cells is more pronounced for lower size ratios (S) compared to higher size ratios. This suggests that the interplay between passive and active cells has a stronger influence on the motility of active cells when their sizes are relatively smaller.

In the subsequent analysis of mean speed for passive cells, we observed a decreasing trend in their velocity with increasing size ratio, as depicted in Figure 3(a) and (b). Since passive cells only experience mechanical forces, this increase in velocity can be attributed to the higher mechanical forces exerted by active cells as the size ratio decreases. As we established earlier, for lower size ratios (S), the active particle’s speed is higher, resulting in stronger mechanical forces acting on the passive cells. Consequently, the passive cells are dispersed at a faster rate. To further support this observation, we plotted the average translational force exerted on the passive cells, as shown in Figure 3(c) and (d). The resulting plots confirm that the average translational force on the passive cells increases with decreasing size ratios, indicating a higher mechanical influence exerted by the active cells. Moreover, for a higher packing fraction of passive cells ϕ*_p_* = 0.5, the mean of the distributions are also higher in all size ratios in comparison to the lower ϕ*_p_* = 0.1. This correlation between size ratio, mean speed of active cells, and the average translational force on passive cells underscores the dynamic interplay between active and passive particles in the system.

**Figure 3:**
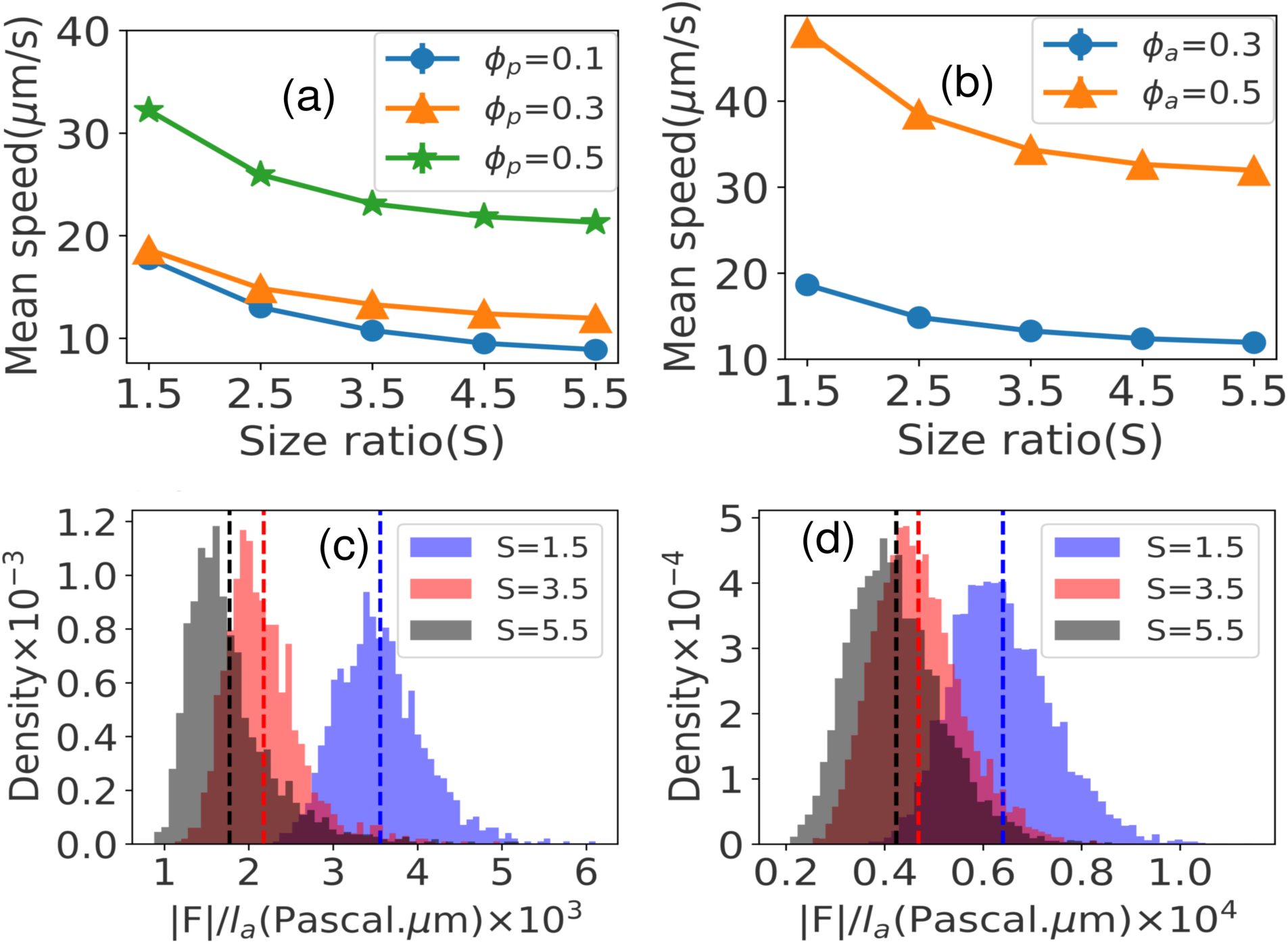
(a) Plot of mean speed of passive cells as a function of size ratios (S) for different ϕ*_p_* and fixed ϕ*_a_* = 0.3 (b) Plot of mean speed of passive cells as a function of size ratios (S) for different ϕ*_a_* and fixed ϕ*_p_* = 0.3. Standard errors are equal to the marker size of the plot.(c) and (d) shows the distribution of modulus value of total force per unit length on a passive cell for fixed ϕ*_p_* = 0.1 and ϕ*_p_* = 0.5 respectively and ϕ*_a_* = 0.3 is fixed for both the plots. The dashed lines shows the mean of the corresponding distributions.

### Velocity autocorrelation

To investigate the dynamical behavior of active cells, we computed the velocity-velocity temporal autocorrelation function, denoted as C(τ) =< ⃗v(x, y, t) · ⃗v(x, y, t + τ) >, where x, y, t, and τ are position, time, and lag time respectively. The angular brackets represent averaging over time and the number of particles in the system. This function quantifies the correlation between an individual cell’s velocity at a given time and its velocity at a later time. By averaging over time and the number of particles, we obtained insights into the decay in autocorrelation of velocity over time. The resulting plots, shown in Figure 4(a) and (b) depicts the temporal velocity autocorrelation for different packing fractions ϕ*_p_* = 0.1 and ϕ*_p_* = 0.5 respectively. Our findings reveal that larger size ratios exhibit slower decay, indicating longer persistence of velocity correlations, while smaller size ratios display faster decorrelation and shorter-lived correlations when the ϕ is smaller. This can be attributed to the weaker alignment interaction among active cells for smaller size ratios, resulting in a quicker randomization of velocity due to collisions at shorter time scales. In contrast, higher size ratios exhibit strong velocity correlations, indicating the formation of organized structures within the colony.

**Figure 4:**
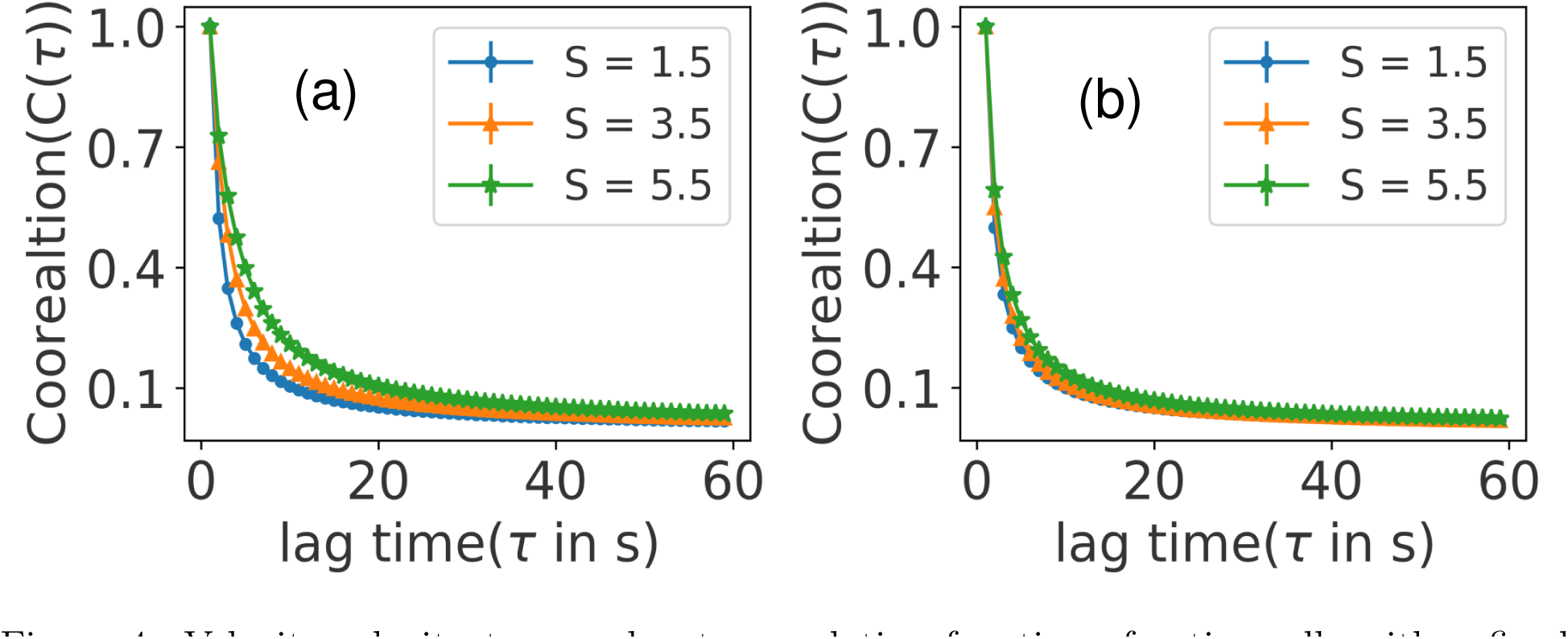
Velocity-velocity temporal auto-correlation function of active cells with a fixed ϕ*_a_* = 0.3 and different values of cell size ratios (S) for: (a) ϕ*_p_* = 0.1 and (b) ϕ*_p_* = 0.5. Standard errors are equal to the marker size of the plot.

However, it is important to note that when the packing fraction is high, as shown in Figure 4(b), the mechanical interactions among the cells become significant. This hinders the alignment of longer cells and causes velocity autocorrelations to rapidly decay, regardless of the size ratios. In such densely packed systems, the intercellular mechanical forces dominate, outweighing the effects of individual cell motility and anisotropy. These findings highlight the complex interplay between cell size ratios, packing fraction, and mechanical interactions in shaping the velocity correlations and emergent structures within the bacterial mixture. Analyzing the velocity autocorrelation function enhances our understanding of the temporal dynamics and interplay of active cells, shedding light on their motion and interactions within the system.

### Mean Squared Displacement (MSD)

To calculate the spatial extent of random motion in our study, we utilized the Mean squared displacement (MSD) for both active and passive cells across various size ratios. The MSD is defined as MSD(τ) = ⟨|x(t + τ) − x(t)|^2^⟩*_t_*, where x(t) represents the position of the cell at time t, τ denotes the lag time, and ⟨⟩*_t_* represents the time average. In general, the MSD can be approximated as MSD(τ) ≈ Dτ *^β^*, where the exponent β identifies the dispersion type: β = 1 for standard diffusion, β < 1 for subdiffusion, and β > 1 for superdiffusion. To characterize the dispersion, we calculated the time-averaged MSD for each cell trajectory and then averaged the results across all trajectories using ensemble averaging. The MSD exhibits two main regimes: the early regime, also known as the ballistic or collision-free regime, and the diffusion regime. Accurately estimating the diffusion coefficient, D, relies on identifying the transition from the ballistic to the diffusion regime. To accomplish this, we employed a fitting algorithm^38^ that utilizes a continuous piecewise function, g(τ, γ, t), with two modes. In this approach MSD is plotted in log-log scale,so that one can distinguish between the two modes more easily.The 2 modes are given as follows:

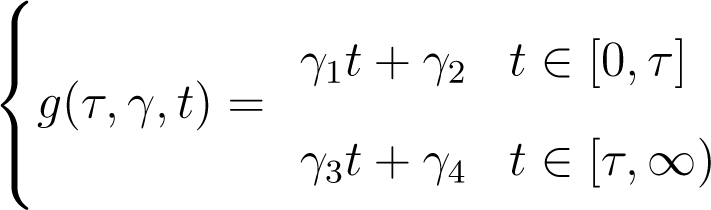

In this algorithm, we treated τ as a parameter for the model and aimed to fit the MSD data with the two pieces of g(τ, γ, t). The accuracy of the fit was assessed using the misfit function, f(t, γ, τ), which is equal to half the sum of squared residuals: 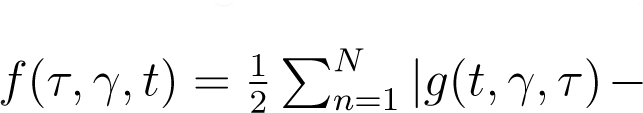 d(t*_n_*)|^2^. Here, d(t*_n_*) represents the discrete data, and N denotes the number of data points. To determine the optimal choice for τ that minimizes the misfit function, we performed an iterative search. By evaluating the misfit function for different values of τ, we could identify the value that yielded the best fit to the MSD data. Through this fitting algorithm, we aimed to accurately extract the breakpoint, or transition point, from the ballistic to the diffusion regime, and obtain reliable diffusion coefficient values for our system.

After determining the optimal τ as the breaking point, we fitted the MSD data using the function g(τ, γ, t). In Figures 5(a),(b) and (c), we present plots of the MSD data of passive cell as a function of time along with the corresponding fitted curves for different size ratios (S). Each of these plots corresponds to a distinct case: Figure 5(a) represents the scenario where ϕ*_a_* > ϕ*_p_*, Figure 5(b) corresponds to ϕ*_a_* = ϕ*_p_*, and Figure 5(c) illustrates ϕ*_a_* < ϕ*_p_*.

**Figure 5:**
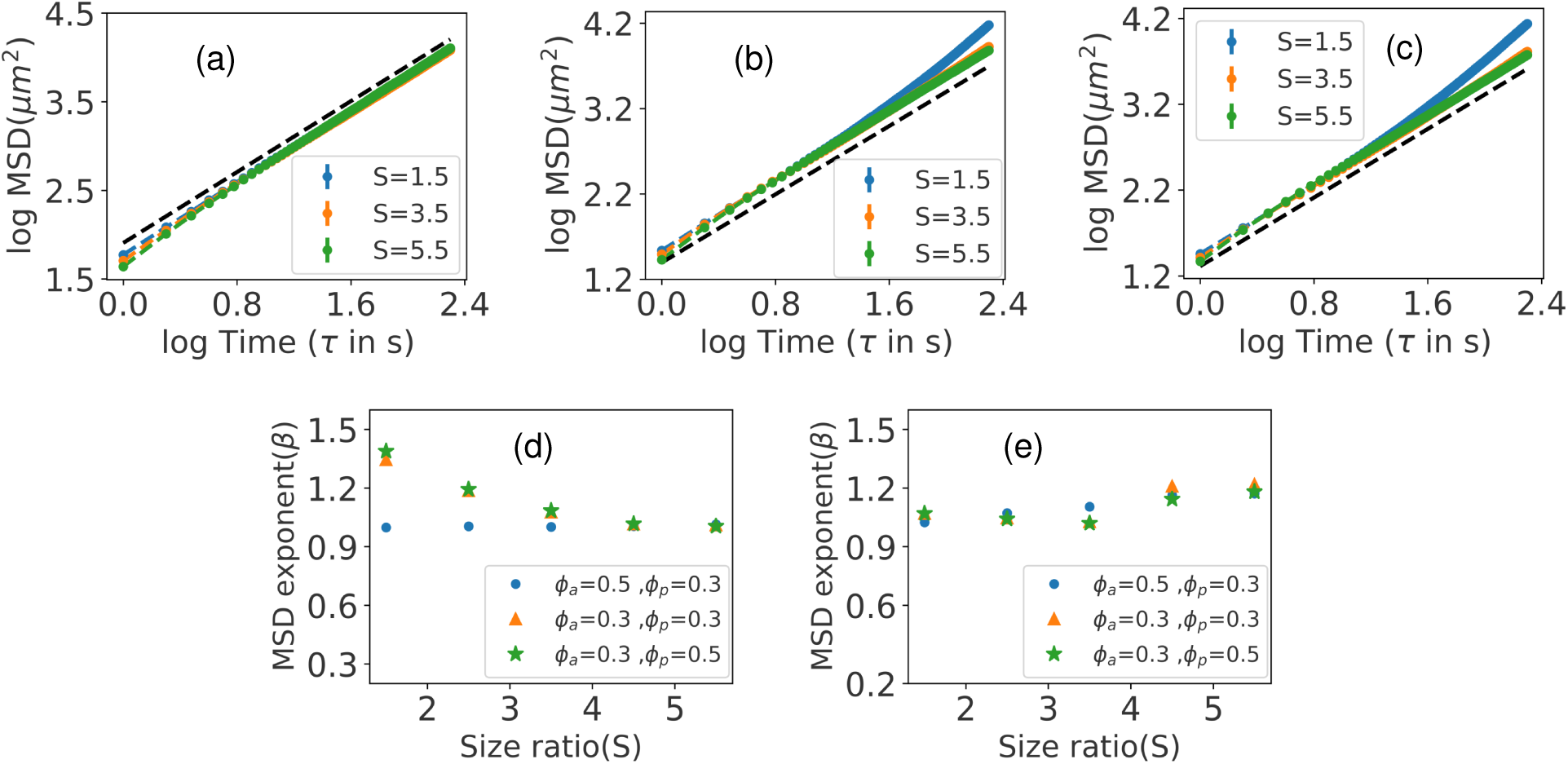
The Mean squared displacement (MSD) of passive cells as a function of lag time for (a) ϕ*_a_*=0.5, ϕ*_p_* =0.3, (b) ϕ*_a_*=0.3, ϕ*_p_*=0.3 and (c) ϕ*_a_*=0.3, ϕ*_p_* =0.5. Standard errors are equal to the marker size of the plot. The dashed black line denotes random Brownian motion with β=1.0. Plot of MSD exponent (β) at (d) diffusion region and (e) ballistic region.

To further quantify this observation, we present the MSD exponent (β) for the ballistic and diffusive regions as a function of size ratio in Figures 5(d) and 5(e), respectively. These plots provide a quantitative measure of the dispersion type and offer insights into the dynamics of the active and passive cells within the system.

i. When (ϕ*_a_* > ϕ*_p_*), indicating a higher area covered by active cells, in diffusion regime the MSD exponent is nearly 1.0. This suggests normal diffusive behavior. Additionally, in the ballistic region, β shows a slight variation with the size ratio (S). Increasing S leads to an increase in β, indicating a more superdiffusive behavior.
ii. When (ϕ*_a_* = ϕ*_p_* = 0.3), meaning equal area coverage by active and passive cells, the behavior of β depends on the size ratio. For smaller S values, β is close to 1.0, indicating diffusive behavior. However, for larger S values, β increases, suggesting superdiffusive behavior. The transition from diffusive to superdiffusive occurs as S increases.
iii. When (ϕ*_a_* < ϕ*_p_*), specifically in the case of ϕ*_a_* = 0.3 and ϕ*_p_* = 0.5, the passive cells exhibit more superdiffusive behavior compared to the previous scenario.

In conclusion, when ϕ*_p_* is smaller than ϕ*_a_* in the diffusion region, β is approximately 1.0, indicating normal diffusive behavior. However, when ϕ*_p_* is equal to or greater than ϕ*_a_*, the passive cells show superdiffusive behavior, especially for smaller size ratios. These findings suggest a relationship between cell activity, area coverage, and the diffusion behavior of the system

A similar analysis was also conducted for the active cells. In the diffusion region, when the exponent β ≈ 1.0, it suggests that most of the active cells exhibit normal diffusive behavior, following a Brownian type of motion across different size ratios and area fractions. However, the behavior is different in the ballistic regime. As shown in Figure 6, it is evident that the value of β increases for larger size ratios, indicating a departure from normal diffusive motion.

**Figure 6:**
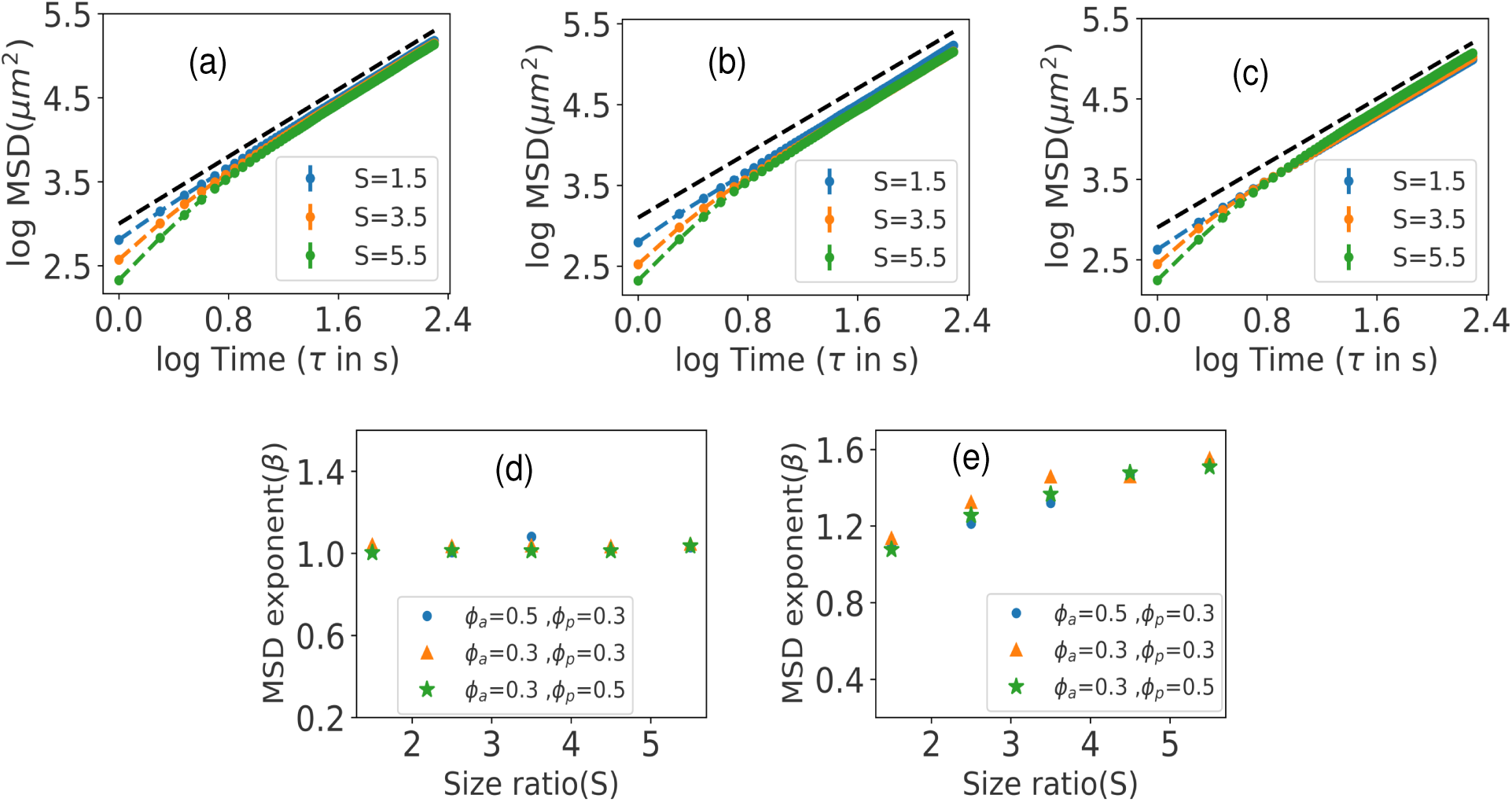
The Mean squared displacement (MSD) of active cells as a function of lag time for (a) ϕ*_a_*=0.5, ϕ*_p_* =0.3, (b) ϕ*_a_*=0.3, ϕ*_p_*=0.3 and (c) ϕ*_a_*=0.3, ϕ*_p_* =0.5. Standard errors are equal to the marker size of the plot. The dashed black line denotes random Brownian motion with β=1.0. Plot of MSD exponent (β) at (d) diffusion region and (e) ballistic region.

### Spatial organization and ordering due to increased size-ratio

Next we focused on computing the radial distribution function, denoted as g(r), to gain deeper insights into the spatial arrangement of active and passive cells within our system. The radial distribution function, also known as the pair correlation function, provides valuable information about the probability distribution of finding a particle at a given radial distance r, from a reference particle’s center. This measurement captures long-range interparticle correlations and reveals the organization of the particles, making it a widely-used tool for characterizing packing structures. In our two-dimensional system, we denote ⟨n⟩g(r)d^2^r as the number of particles in d^2^**r**, where r = |⃗r_1_ − ⃗r_2_|. Here, ⃗r_1_ represents the position of the reference particle, and ⃗r_2_ represents the position of the neighboring particle. To comprehensively understand the structural characteristics of our system, we define two types of radial distribution functions: g*_pp_*(r) for passive cell pairs and g*_aa_*(r) for active cell pairings.

In the subsequent sections, we thoroughly investigate these distribution functions as a function of the passive cell area fraction (ϕ*_p_*), the active cell area fraction (ϕ*_a_*), and the particle size ratio (S). To initiate our investigation, we explore the influence of the size ratio (S) on the radial distribution function of passive particles, g*_pp_*(r), while keeping ϕ*_a_* fixed at 0.3 (Figure 7(a), (b) and (c)). We conduct this analysis across three distinct ϕ*_p_* values: 0.1, 0.3 and 0.5. We observe several significant features regarding the behavior of the passive cells:

**Figure 7:**
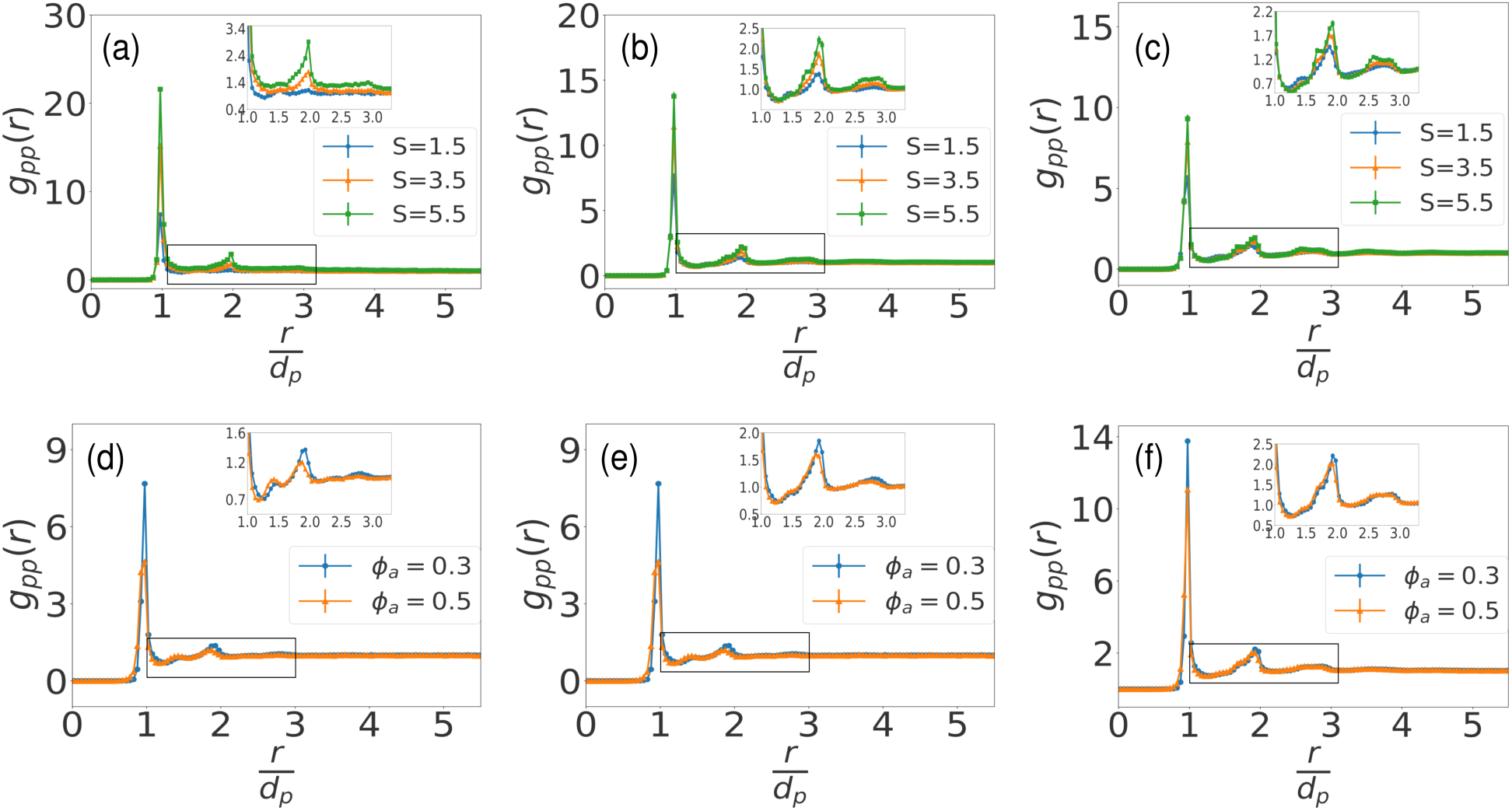
Radial distribution function or pair-correlation function (g*_pp_*(r)) for passive cells for three different size ratios and a fixed active cell area fraction (ϕ*_a_*=0.3): (a) ϕ*_p_*=0.1, (b) ϕ*_p_*=0.3 and (c) ϕ*_p_*=0.5. Plots of g*_pp_*(r) at a fixed ϕ*_p_* = 0.3 and two different ϕ*_a_*s for (d) S=1.5, (e) S=3.5 and (f) S=5.5. Peaks are more visible in the enlarged image in the inset.

(i) In all cases depicted in Figure 7(a), (b), and (c), we observe a peak at r = d*_p_*, which corresponds to the diameter of the passive particle. This peak indicates the presence of nearest-neighbor interactions between passive particles. (ii) Additionally, we observe the emergence of a smaller peak at r = 1.9d*_p_*, which becomes more pronounced as the size ratio increases. This peak suggests that the passive particles are arranged in a hexagonally close-packed configuration within the cluster. This arrangement holds true for all investigated cases. (iii) For higher values of ϕ*_p_* (0.3 and 0.5), as shown in Figure 7(b) and (c), a distinct peak is observed at a slightly larger distance, r = 2.7d*_p_*. This peak indicates the formation of an ordered aggregate with well-defined layering of the coordination shell. However, this peak is absent for a smaller area fraction of passive cells (ϕ*_p_* = 0.1). These observations from the g*_pp_*(r) plots suggest that the effective attractive interaction between passive particles induced by the active particles becomes more significant for larger size ratios. Notably, for a size ratio of 5.5, the peaks in the radial distribution function are the most prominent, indicating the formation of robust clustering among the passive particles.

Next, in order to gain further insights into the effective interaction between passive cells, we conducted an analysis for two different values of ϕ*_a_* while keeping ϕ*_p_* constant at 0.3 (Figure 7(d), (e) and (f)). (i) We observe that the first peak in g*_pp_*(r) still occurs at r = d*_p_* for all size ratios. However, it is evident that for ϕ*_a_* = 0.3, all the peaks in g*_pp_*(r) are larger compared to ϕ*_a_* = 0.5. This implies that for a lower area fraction of active cells, the passive cells exhibit a more pronounced structural arrangement. (ii) For a small size ratio (S = 1.5) and both values of ϕ*_a_*, a small peak is observed at r = 1.3d*_p_*, indicating the presence of a second layer of passive particles. Furthermore, a relatively larger third peak emerges at r = 1.9d*_p_*. However, as the size ratio increases, the second peak at r = 1.3d*_p_* diminishes, and g*_pp_*(r) exhibits a more prominent peak at r = 1.9d*_p_*. This suggests the formation of dense clusters of passive cells with increasing size ratio.

The overall structure of the binary mixture of morphologically distinct cells can be better understood by considering the tendency of active particles to aggregate. To investigate this, we plotted the radial distribution function for the active particles as a function of the size ratio (S), as shown in Figure 8. It is evident that a peak appears close to r = d*_a_*, indicating the presence of inter particle interactions. Notably, as the size ratio S increases, this peak becomes more pronounced, broadens, and shifts towards r = 0.5d*_a_*. It suggests that as we increase the size ratio while maintaining a fixed area fraction of particles, the active particles become more closely packed together.

**Figure 8:**
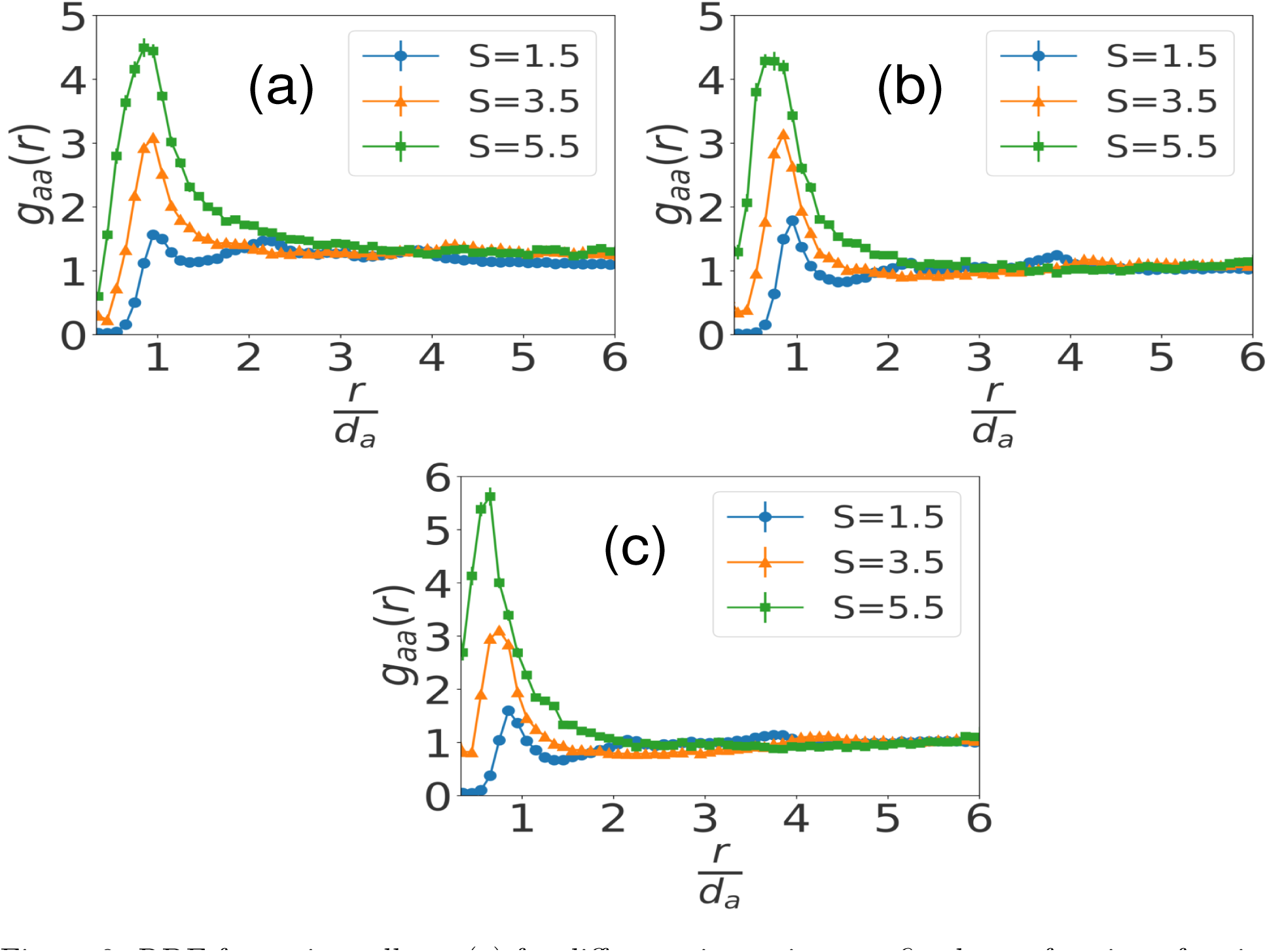
RDF for active cells g*_aa_*(r) for different size ratios at a fixed area fraction of active cells (ϕ*_a_*=0.3) for varying packing fraction of passive cells: (a) ϕ*_p_*=0.1, (b) ϕ*_p_*=0.3 and (c) ϕ*_p_*=0.5.

Furthermore, for a specific size ratio of S= 1.5, additional small peaks are observed in the radial distribution function of active particles. These peaks occur at r = 2.1d*_a_* and r = 3.9d*_a_*, indicating the formation of structures at those distances. However, as the size ratio (S) increases, these secondary peaks become narrower and the peak at r = d*_a_* becomes broader. This behavior suggests a compression or squeezing of the structures at intermediate distances, while the alignment of active cells becomes more prominent. In what follows we will now characterize the emerging ordering in terms of aggregation and clustering as evident within the binary system.

### Clustering and cluster size distribution

Motivated by previous studies,^27, 30, 39^ we conducted an analysis of the cluster size distribution (CSD) for both passive and active bacterial cells, considering different size ratios and area fractions. The CSD, denoted as p(n), represents the probability of having n number of cells in a cluster. In our study, clusters were identified based on a distance cutoff criterion. Specifically, we classified two passive cells as belonging to the same cluster if their separation distance was less than or equal to 1.1 times the diameter of a passive cell (d*_p_*). This cutoff value was chosen based on the first peak observed in the radial distribution function.

To initiate our analysis, we systematically varied the area fraction of passive particles (ϕ*_p_*), area fraction of active particles (ϕ*_a_*) and size ratio (S) one by one by keeping the other quatities fixed. In Figure 9(a)-(d), we illustrated the cluster size distribution (CSD) for passive cells. Based on the results, we can make the following observations:

i. For ϕ*_p_* = 0.1 and lower S such as 1.5 and 2.5, the CSD follows an exponential form, characterized by exp(−n/n_0_). This suggests weak clustering at these size ratios. However, for S > 2.5, the CSD exhibits a power-law decay with an exponential cutoff at larger cluster sizes (n = n_0_)[Figure 9(a)]. A good fit to our data is given by p(n)/p(1) ≈ ^1^ exp(−n/n_0_).

**Figure 9:**
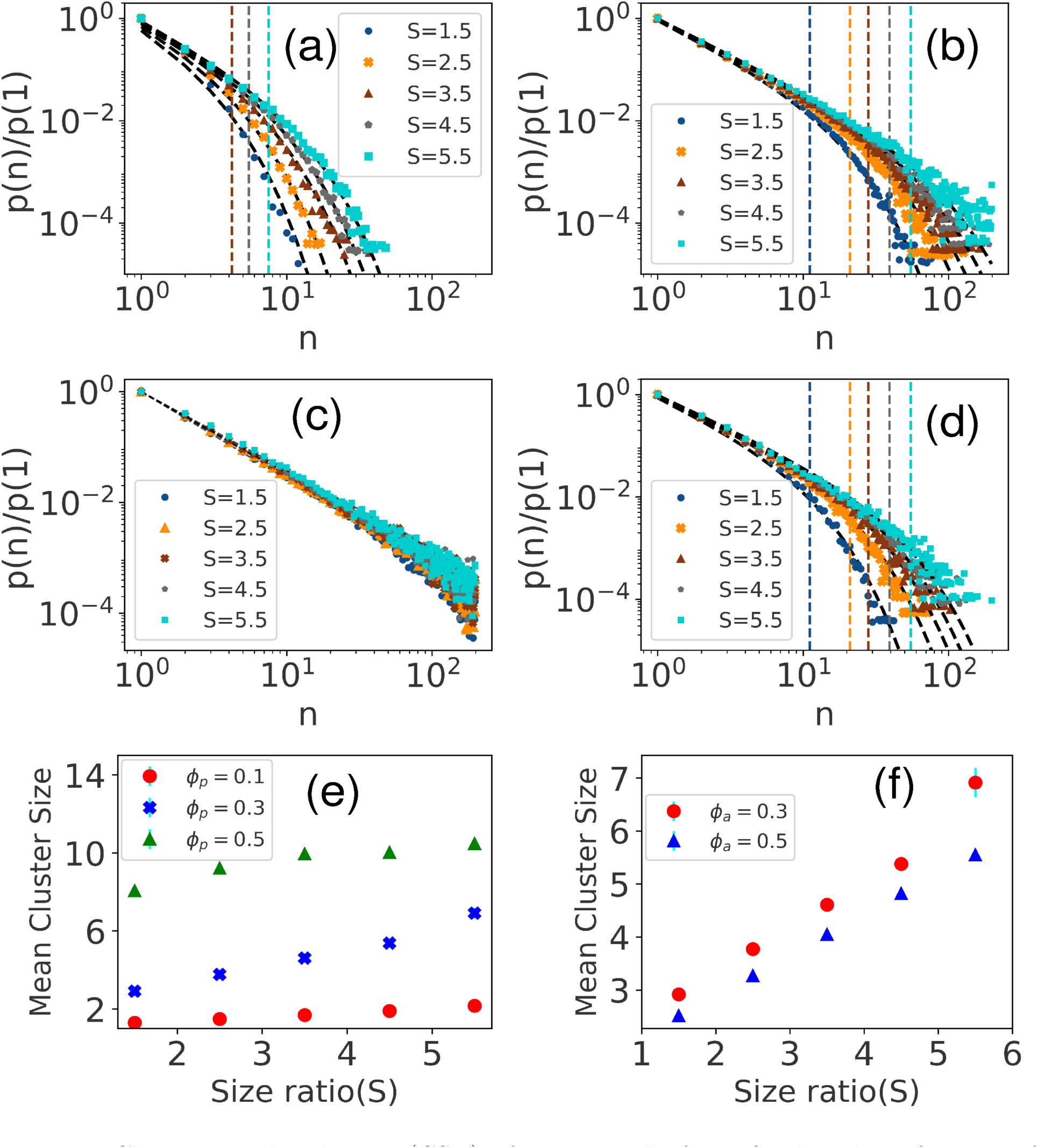
Cluster size distribution (CSD) of passive cells for a fixed packing fraction of active cells ϕ*_a_*=0.3 for different size ratio S. (a) ϕ*_p_*=0.1, (b) ϕ*_p_*=0.3 and (c) ϕ*_p_*=0.5. (d) CSD plots of passive cells when ϕ*_a_*=0.5 and ϕ*_p_*=0.3. The dashed lines signifies the cut-off n_0_ for different S and to differentiate the dashed line with its corresponding fit line has the same color. Plot of average cluster size of passive cells as a function of size ratios for (e) different ϕ*_p_* at fixed ϕ*_a_*=0.3 and (f) for different ϕ*_a_* at fixed ϕ*_p_*=0.3.

This indicates the presence of clustering phenomena at larger size ratios.

(ii) Moving to intermediate ϕ*_p_* = 0.3 and varying S, the CSD plot can still be described by p(n)/p(1) ≈ ^1^ exp(−n/n_0_). However, it is worth noting that the distribution shifts to the right as the value of the exponential cutoff (n_0_) increases with changing size ratios [Figure 9(b)]. This implies that the occurrence of larger clusters becomes more probable. To illustrate this, we have included vertical dashed lines representing the cutoff for each size ratio.
(iii) At a higher ϕ*_p_* = 0.5, the CSD plot follows a nearly normal power-law decay of the form p(n)/p(1) ≈ ^1^

[Figure 9(c)]. This suggests that the system is approaching phase separation. Increasing the size ratio (S) also leads to a decrease in the exponent of the power law. For example, at S = 1.5, α = 1.58, while at S = 5.5, α = 1.42. This indicates that increasing the size ratio enhances clustering.

To investigate the effect of the number of active cells, we also analyzed the CSD plot for a higher area fraction of active particles (ϕ*_a_*=0.5) in Figure 9(d). Similar to the earlier cases, here also the CSD is well described by a power law with an exponential cutoff. To further understand the dynamics of clustering, we calculated the average cluster size of passive cells as a function of the size ratio (S) using the formula ⟨n⟩ = n · p(n) for various area fractions of active cells (ϕ*_a_*) and passive cells (ϕ*_p_*). Two plots were created to represent the average cluster sizes for passive cells (Figure 9(e) and (f)). Both figures clearly demonstrate that increasing the size ratio or the length of elongated active cells leads to larger cluster sizes. Moreover, from the mean cluster size plot in Figure 9(e), it can be observed that for a fixed size ratio, increasing ϕ*_p_* results in larger cluster sizes. But, in Figure 9(f), for a particular S and ϕ*_p_*, increasing the area fraction of active particles (ϕ*_a_*) also decreases the cluster size. This implies that, for the same number of passive cells, the presence of active cells has an impact on the cluster size of the passive cells. These findings are further supported by the radial distribution function (RDF) plot in Figure 7, where the peaks of g*_pp_* are notably lower when ϕ*_a_* is greater.

In Figure 10, we present the corresponding plots of the cluster size distribution (CSD) of active cells, considering different size ratios and area fractions. Based on the CSD results, we observe that for a higher ϕ*_a_* = 0.5 (Figure 10(b)), apparently as S increases, the distribution is slightly shifted towards the right. This indicates that when the size ratio is larger, elongated active cells tend to become more clustered. However, it is worth noting that we couldn’t find a specific functional form to fit these CSD plots for the active cells. In summary, the plots of the cluster size distribution for the active cells demonstrate that for a higher area fraction of active particles, increasing the size ratio results in a shift towards more clustering of elongated active cells. However, further analysis is required to determine a specific functional form to describe these distributions accurately.

**Figure 10:**
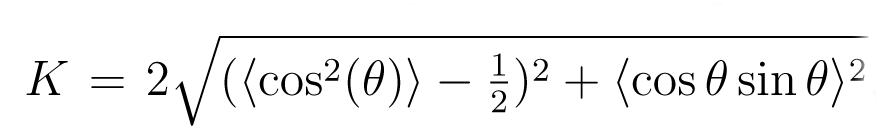
Cluster size distribution (CSD) for active cells for different size ratios for fixed ϕ*_p_*=0.3 and for (a) ϕ*_a_*=0.3 and (b) ϕ*_a_*=0.5.

### Nematic phase transition within the mixture

In order to gain further insights into the local orientational order within the system, we determined the scalar nematic order parameter (K) to assess the degree of alignment in the nematic phase. The nematic phase is characterized by the alignment of objects or particles in the same direction. Inspired by previous studies,^40, 41^ we employed the scalar nematic order parameter to quantify the alignment of rod-shaped active cells in our system. The expression for the 2D nematic order parameter^42^ is given by 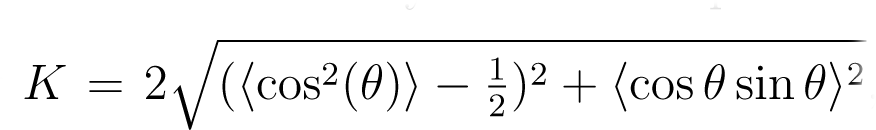, where θ represents the angle between the long axis of neighboring particles. The angle brackets indicate averaging over the entire system. A value of K = 1 corresponds to a perfectly aligned phase, while K = 0 indicates an isotropic phase. We plotted the distribution of K for the active cells across different S values to analyze the alignment characteristics within the system.

In the probability distribution, P(K), we have ensured normalization such that ^J^ P(K)dK = 1. For neighbor searching, a cutoff distance of r*_cut_* = 10.0/d*_a_* has been chosen for each particle, where d*_a_* represents the diameter of an active cell. Figure 11 depicts the distribution of P(K), revealing that as the size ratio increases, the distribution becomes broader, indicating larger values of K. This suggests that an increasing size ratio induces ordering among the active cells, leading to a higher nematic order parameter.

**Figure 11:**
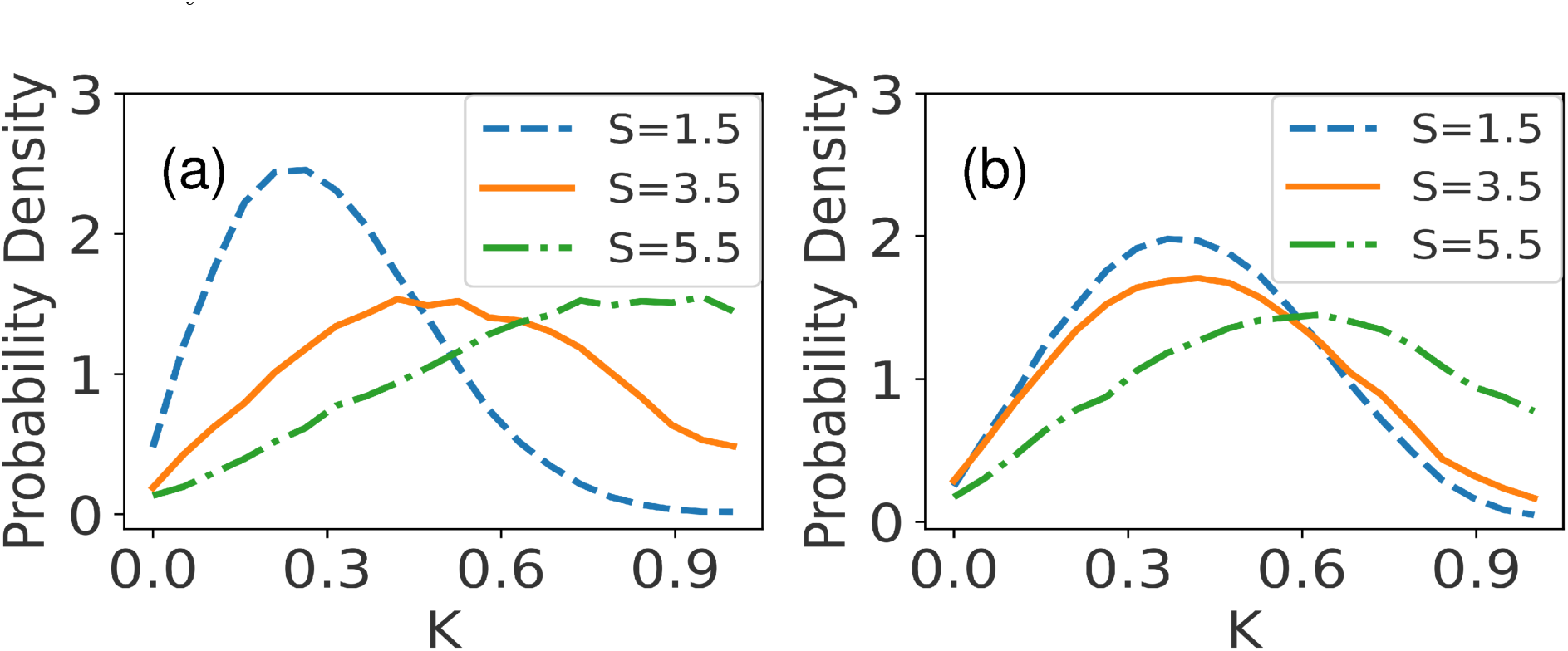
Distribution of the scalar nematic order parameter K of active cells at a fixed ϕ*_a_* = 0.3 and for different values of S for (a) ϕ*_p_* = 0.1, and (b) ϕ*_p_* = 0.5.

Furthermore, we observed a significant influence of the passive cell fraction on the response towards nematicity, as depicted in Figure 11(a) and (b). For a lower size ratio of S = 1.5, an increase in ϕ*_p_* results in a slight enhancement of the maximum K value. In contrast, for S = 5.5, an increase in ϕ*_p_* causes a shift towards the left, resulting in a lowering of nematic order. This highlights that the passive cell fraction plays a crucial role in determining the response towards nematicity, and its effect is dependent on the size ratio.

## Concluding remarks

Microbial species often coexist in complex natural systems, and understanding their dynamics is crucial for various applications. In this study, we explore the dynamics of a mixed bacterial population consisting of two species with distinct morphology and motility properties. We aim to understand how the motility and anisotropy of one species can drive dispersal, aggregate formation, and clustering of spherical nonmotile species. By developing an agent-based model and employing computer simulations and systematic analysis, we investigate the spatiotemporal dynamics and emergent ordering in such systems. By simulating the system for different size ratios at a fixed packing fraction, we observed distinct patterns of dispersal, aggregate formation, and clustering. The motility and anisotropy of one species significantly influenced the distribution and spatial organization of the nonmotile spherical species. Furthermore, varying the packing fraction for a fixed size ratio revealed additional insights into the system’s behavior, demonstrating the importance of interplay between surface fractions and species characteristics.

Specifically, we observe nonmonotonic changes in the mean speed of passive cells with respect to their packing fraction, especially for smaller size ratios, resulting in superdiffusive behavior. Active cells exhibited normal diffusive behavior in the diffusion regime, while for higher size ratios, they displayed superdiffusivity in the ballistic regime. Moreover, the increase in size ratio leads to the clustering of passive cells, while active cells become more aligned and closely packed in the presence of higher surface fractions of passive cells. Additionally, we have identified that the size ratio and packing fraction of passive cells influence the nematic order and ordering of active cells. Our results highlight the intricate relationship between motility, anisotropy, and emergent ordering in mixed bacterial communities. The observed dynamics provide valuable insights into the mechanisms driving dispersal, aggregate formation, and clustering.

In conclusion, our study elucidates the spatiotemporal dynamics and emergent ordering in a mixed bacterial population comprising morphologically distinct species. Our results demonstrate the importance of surface fraction and size ratio in shaping the system’s organization. These findings contribute to the broader understanding of microbial community dynamics and have practical implications for various applications involving bacterial populations. Future research can build upon these insights to develop strategies for manipulating and controlling microbial communities for desired outcomes.

## Acknowledgement

We greatly acknowledge Indian Institute of Science Education and Research Thiruvananthapuram (IISERTVM), India for providing the support of computing resources. We also acknowledge the support for high-performance computing time at the Padmanabha cluster, IISERTVM, India. P.G. acknowledges Start-up Research Grant (SRG/2022/000043) by Science and Engineering Research Board (SERB), India.

## Supporting Information Available

The supporting information (SI) contains video files mentioned in the main text.

